# The virucidal effects of 405 nm visible light on SARS-CoV-2 and influenza A virus

**DOI:** 10.1101/2021.03.14.435337

**Authors:** Raveen Rathnasinghe, Sonia Jangra, Lisa Miorin, Michael Schotsasert, Clifford Yahnke, Adolfo Garcίa-Sastre

## Abstract

Germicidal potential of specific wavelengths within the electromagnetic spectrum is an area of growing interest. While ultra-violet (UV) based technologies have shown satisfactory virucidal potential, the photo-toxicity in humans coupled with UV associated polymer degradation limit its use in occupied spaces. Alternatively, longer wavelengths with less irradiation energy such as visible light (405 nm) have largely been explored in the context of bactericidal and fungicidal applications. Such studies indicated that 405 nm mediated inactivation is caused by the absorbance of porphyrins within the organism creating reactive oxygen species which result in free radical damage to its DNA and disruption of cellular functions. The virucidal potential of visible-light based technologies has been largely unexplored and speculated to be ineffective given the lack of porphyrins in viruses. The current study demonstrated increased susceptibility of lipid-enveloped respiratory pathogens of importance such as SARS-CoV-2 (causative agent of COVID-19) as well as the influenza A virus to 405nm, visible light in the absence of exogenous photosensitizers indicating a potential porphyrin-independent alternative mechanism of visible light mediated viral inactivation. These results were obtained using less than expected irradiance levels which are generally safe for humans and commercially achievable. Our results support further exploration of the use of visible light technology for the application of continuous decontamination in occupied areas within hospitals and/or infectious disease laboratories, specifically for the inactivation of respiratory pathogens such as SARS-CoV-2 and Influenza A.

## Introduction

The severe-acute respiratory syndrome corona virus 2 (SARS-CoV-2), the causative agent of the COVID-19 pandemic, is a member of the beta-coronavirus family and it emerged at the end of 2019 in the Hubei province in Wuhan China^1^. By late February 2021, more than 112 million cases had been reported while accounting for approximately 2.5 million deaths, underscoring the rapid dissemination of the virus on a global scale^2^. As a complement to standard precautions such as handwashing, masking, surface disinfection, and social distancing, other enhancements to enclosed spaces such as improved ventilation and whole-room disinfection are being considered by segments beyond acute healthcare such as retail, dining, and transportation^3^.

Initial guidance from health authorities such as the CDC and WHO on environmental transmission focused on contaminated surfaces as fomites^4^. Data pertaining to the survival of SARS-CoV-2 and other related coronaviruses to date has indicated that virions are able to persist on fomites composed of plastic^5^, wood^6^, paper^5^, metal^7^ and glass^8^ potentially up to nine days. Recent studies have suggested that SARS-CoV-2 may also remain viable approximately at least three days in such surfaces and another two studies showed that at room temperature (20-25°C), a 14-day time-period was required to see a 4.5-5 Log_10_ of the virus^9, 10^.

Since the start of the pandemic, transmission of the virus by respiratory droplets and aerosols has become an accepted method of transmission although the relative impact of each mode of transmission is the subject of much debate. Nevertheless, enclosed spaces with groups of people exercising or singing have been associated with increased transmission. The half-life survival of SARS-CoV-2 in this type of environment has been estimated between 1-2 hours^6, 11, 12^.

Taking this information into consideration, several methods have been evaluated to effectively inactivate SARS-CoV-2. Chemical methods, which focus on surface disinfection, utilize 70% alcohol and bleach and their benefits are well established. These methods are also episodic (or non-continuous) meaning that in-between applications, the environment is not being treated^13^.

In addition to chemicals, one of the most utilized methods for whole-room disinfection is germicidal ultra-violet C (UVC; ~254 nm)^14^. This technology is well established^15^ and has been shown to inactivate a range of pathogens including bacteria^16^, fungi^17^ and viruses^18^. The mechanism of action of UVC is photodimerization of genetic material such as RNA (relevant for SARS-CoV2 and IAV) and DNA (relevant for DNA viruses and bacterial pathogens, among others)^19^. Unfortunately, this effect has been associated with deleterious effects in exposed humans such as photokeratoconunctivitis in eyes and photodermatitis in skin^20^. For these reasons, UVC irradiation requires safety precautions and cannot be used to decontaminate fomites and high contact areas in the presence of humans^21^.

Germicidal properties of violet-blue visible light (380-500 nm), especially within the range of 405 to 450 nm wavelengths have been appreciated as an alternative to UVC irradiation in whole-room disinfection scenarios where it has shown reduction of bacteria^22, 23^ in occupied rooms and reductions in surgical site infections^24^. Although 405 nm or closely related wavelengths have been shown to be less germicidal than UVC, its inactivation potential has been assessed in pathogenic bacteria such as *Listeria* spp and *Clostridium* spp^24, 25^, and in fungal species such as *Saccharomyces* spp and *Candida* spp^26^. It is thought that the underlying mechanism of blue-light mediated inactivation is associated with absorption of light via photosensitizers such as porphyrins which results in the release of reactive oxygen species (ROS) ^27, 28^. The emergence of ROS is associated with direct damage to biomolecules such as proteins, lipids and nucleic acids which are essential constituents of bacteria, fungi and viruses. Further studies have shown that ROS can also lead to the loss of cell membrane permeability mediated by lipid oxidation^29^. Given the lack of endogenous photosensitizers such as porphyrins in virions, efficient decontamination of viruses (both enveloped and non-enveloped) may require the addition of exogenous photosensitizers^23^. With the use of media suspensions containing both endogenous and/or exogenous photosensitizers, inactivation of viruses such as feline calcivirus (FCV)^30^, viral hemorrhagic septicemia virus (VHSV)^31^ and murine norovirus-1^32^ has demonstrated the virucidal potency of 405 nm visible light using porphyrin rich media such as saliva, blood, etc. (to create the ROS required for inactivation) using unsafe, commercially impractical irradiance levels. This highlights the importance of answering the basic scientific questions related to viral inactivation within the context of the applied science required for clinical application. Our study specifically focused on three questions: 1) does the same wavelength of light (405nm) that inactivates bacteria also inactivate enveloped viruses, 2) is a porphyrin rich medium required for this inactivation, and 3) can this inactivation be achieved using safe, commercially practical irradiance levels?

In the current study, we show the impact of 405 nm irradiation on inactivation of SARS-CoV-2 and influenza A H1N1 viruses without the use of photosensitizers making it directly relevant to the clinical environment. We show this using a commercially available visible light disinfection system ensuring that the irradiance used is both safe and practically achievable in a clinical setting supporting the possible use of 405 nm irradiation as a tool to confer continuous decontamination of respiratory pathogens such as SARS-CoV-2 and influenza A viruses. We further show the increased susceptibility of lipid-enveloped viruses for irradiation in comparison to non-enveloped viruses, further characterizing the virucidal effects of visible light.

## Materials and methods

### 405 nm Exposure System

The visible light disinfection product used in this study was a commercially available 6” LED downlight (Indigo-Clean, Kenall, Kenosha, WI) to allow for use within a BSL-3 level containment hood. Within the hood, the distance between the face of the fixture and the sample was 10”- much less than the normal 1.5 m used in normal, whole-room disinfection applications. The output of the fixture was modified electronically during its manufacture to match this difference and ensure that the measurements would represent the performance of the device in actual use. This test setup is shown in a BSL-2 hood in Figure 1 below. Note the spectroradiometer and the bottom portion of the rig (~6”) used for calibration were removed during the actual study.

**Figure 1.**
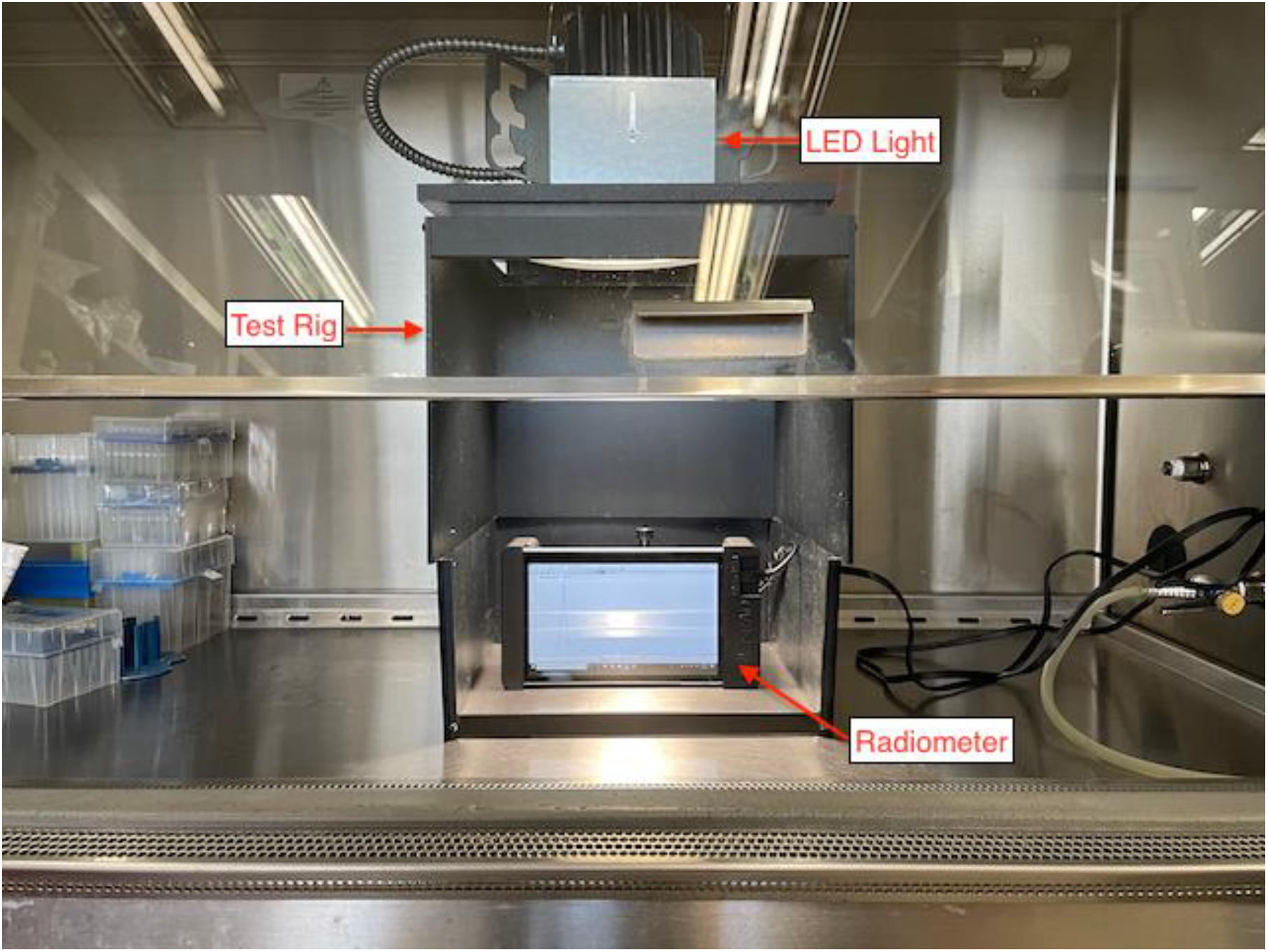
Test setup shown with spectroradiometer and extension used for calibration.

Controls were placed outside the test rig but within the BSL hood as shown in Figure 2. Note that this picture contains the bottom portion of the test rig to highlight the position of the radiometer

**Figure 2.**
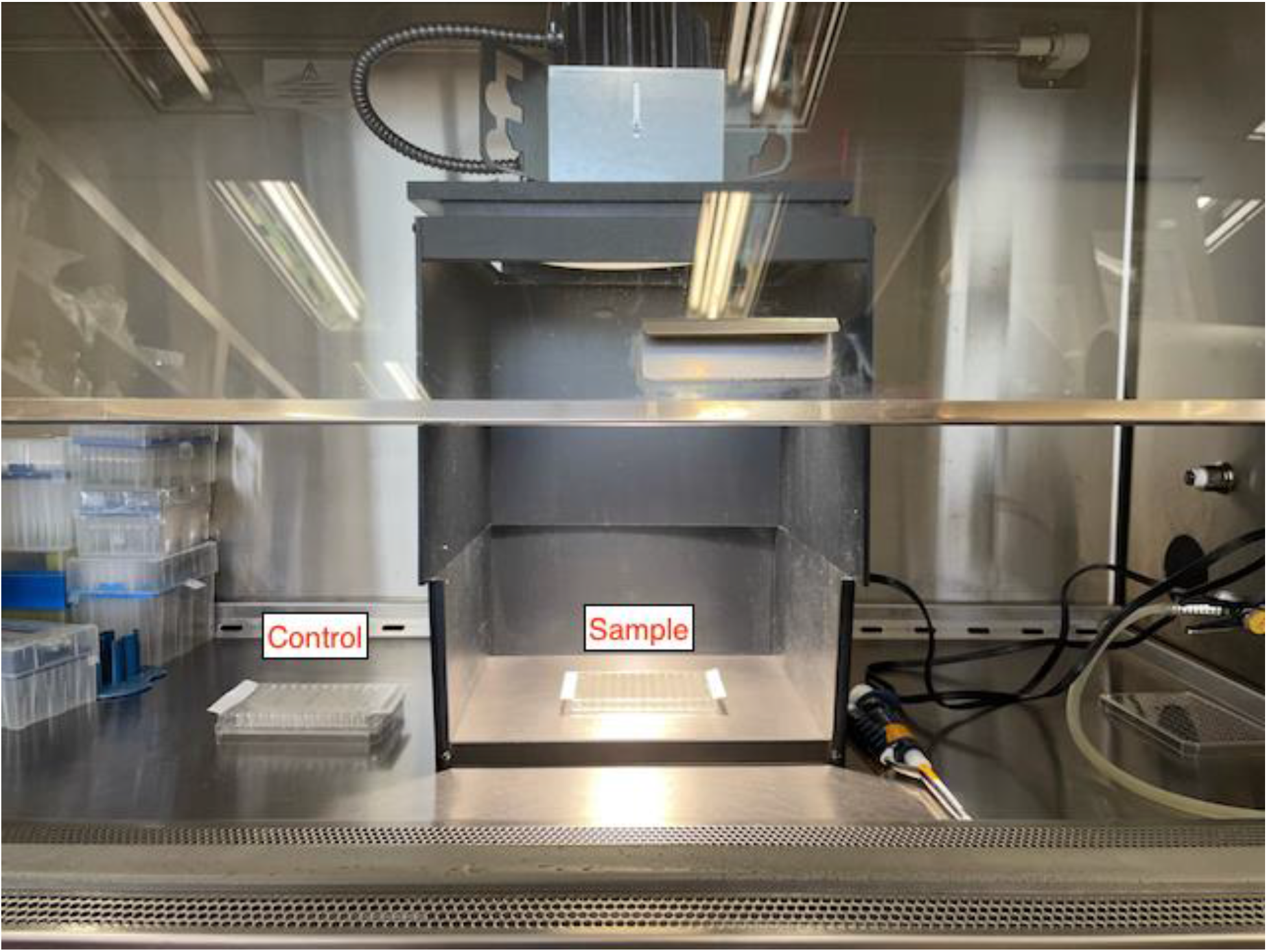
Test setup showing the placement of the control and sample for irradiation. Note that the bottom portion of the test rig was removed during the actual experiment. This ensures the 10” distance used in the study.

For the range of output used in this study, multiple discrete levels were created using pulse width modulation within the LED driver itself. These levels were made to be individually selectable using a simple knob on the attached control module.

As expected, the amount of visible light within the 400nm-420nm bandwidth, measured in mWcm^−2^, is a measurement of the “dose” delivered to the target organism and is used to quantify this relationship similar to that used in UV disinfection applications.

To fully examine this effect, a range of irradiance values were used representing actual product deployment conditions in occupied rooms. The lowest value (0.035 mWcm^−2^) represents a single-mode, lower wattage used in general lighting applications while the highest value (0.6 mWcm^−2^) represents a dual-mode, higher wattage used in critical care applications such as an operating room.

The device was placed in a rig to ensure a consistent distance (10”) between the fixture and the samples. The output of the fixture in the test rig was measured using a Stellar-RAD Radiometer from StellarNet configured to make wavelength and irradiance measurements from 350nm-1100nm with < 1nm spectral bandwidth using a NIST traceable calibration. To ensure that the regular white light portion of the illumination (which is non-disinfecting) was not measured, the measurement was electronically limited to a 1nm bandwidth over the 400nm-420nm range. The normalized spectral profile is shown in Fig. 3 below. The absolute value of the measurement was determined using a NIST traceable calibration as previously described.

**Figure 3.**
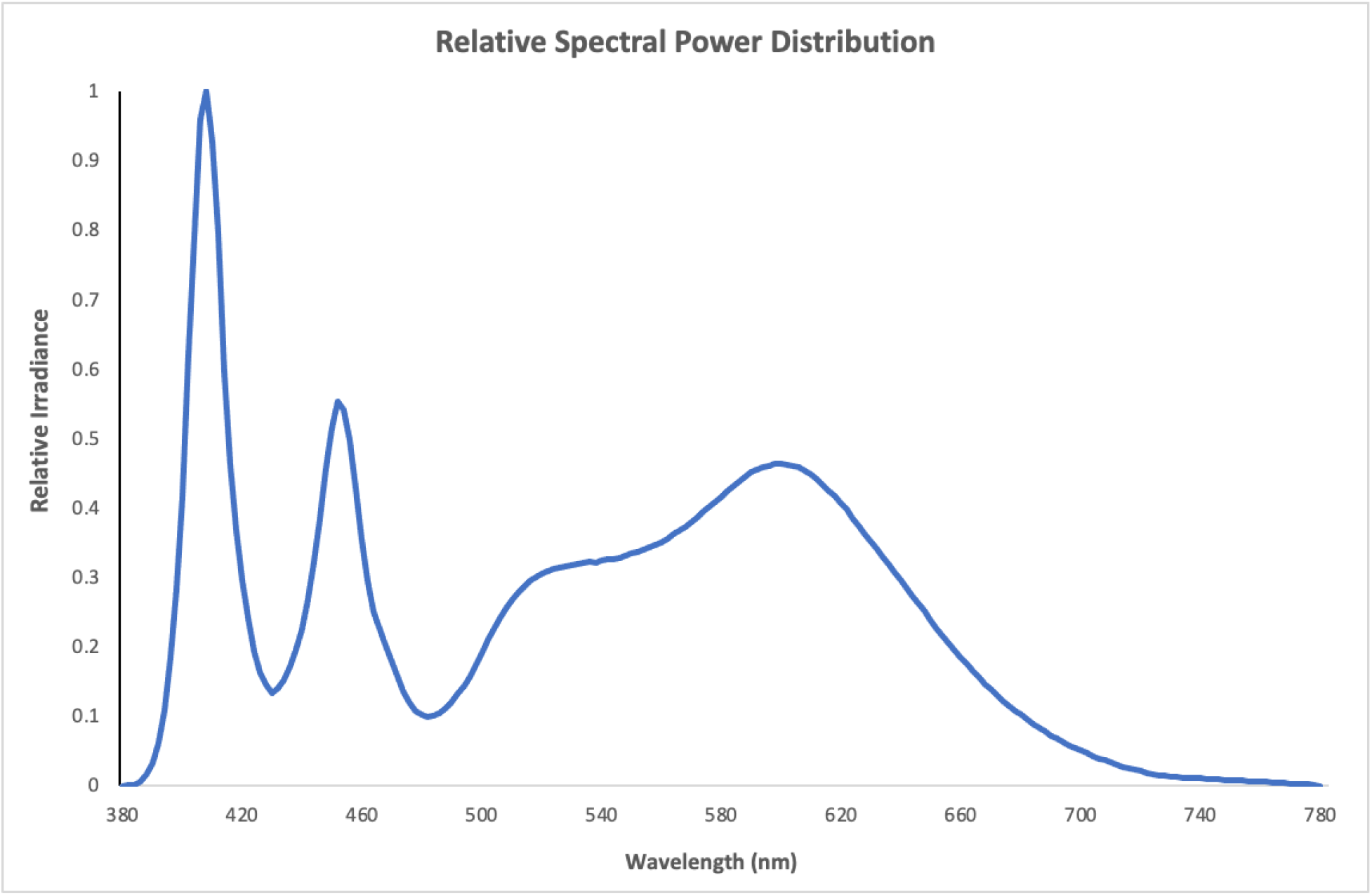
Normalized spectral power distribution showing peak irradiance at 405nm.

Samples were irradiated with the range of wavelengths depicted in Figure 3. This was deliberately done for two reasons: 1) prior work had shown that visible light disinfection was primarily active at 405nm^33^ and 2) to emphasize the applied science associated with actual clinical use where a virus in the environment could be exposed to both 405nm and regular white light in an occupied room.

To isolate the contribution of 405nm light, the control samples were placed outside the field of irradiation created by the disinfection product but within the biosafety hood. Accordingly, these samples were exposed to the overhead lights within the room which contained virtually no 405nm light (< 0.001 mWcm^−2^).

In any assessment of viral inactivation, thermal denaturing of the organism is a concern. Older lighting technologies such as incandescent sources heat a resistive element and were widely used in a variety of applications. This creates heat at both the source and to objects within its field. Fortunately, the disinfecting light (sample) and the overhead lights in the room (control) did not use this technology and therefore contain no infrared emissions (> 800 nm) a commonly known benefit associated with LED lighting. As a confirmation, the temperature beneath the disinfecting light was measured using a commercially available thermocouple during a 24-hour period. During this time, the temperature at the sample position was constant at 20°C +/− 0.5°C. even at the highest disinfecting power (0.6 mWcm^−2^).

For reference, two spectral profiles, one for traditional fluorescent lighting and the other for standard LED lighting are provided in Figure 4 below. While both types of lighting are commonly used in the overhead lights within buildings, the lights in the BSL-3 laboratory were traditional fluorescent. As shown in Figure 2, the controls were exposed only to traditional fluorescent lighting with a negligible amount of disinfecting light (< 0.001 mWcm^−2^) between 400nm and 420nm. Due to the inherent differences between fluorescent and LED lighting, the standard LED spectra has a small, but measurable amount of disinfecting light (0.006 mWcm^−2^) between 400nm and 420nm. As will be later shown, this amount of light can have a measurable disinfecting effect.

**Figure 4.**
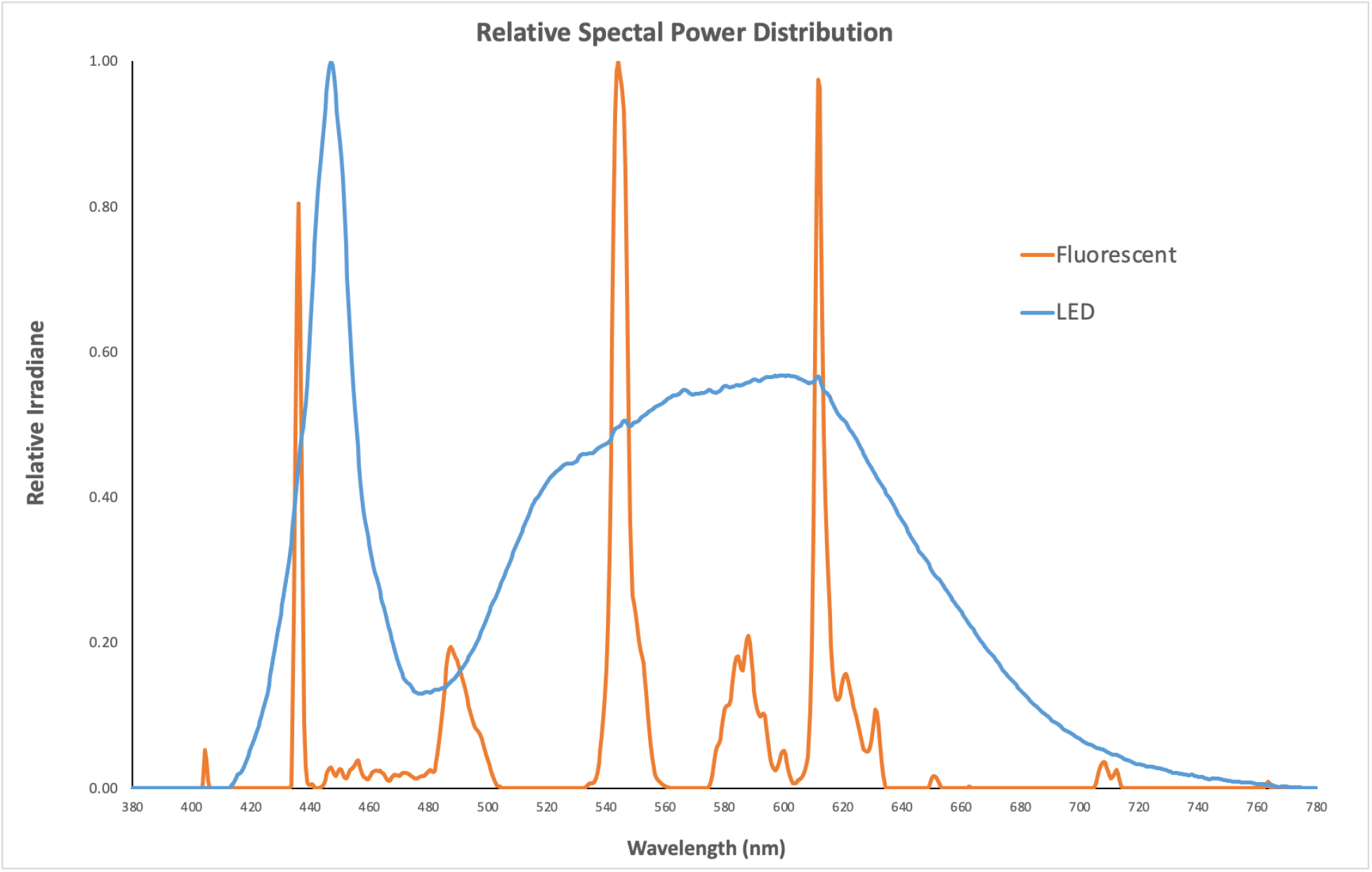
Normalized spectral power distribution for the fluorescent control light (non-disinfecting) and standard LED light (without 405nm) used in the study. Each spectrum is normalized relative to its own peak value.

### Cells and viruses

Vero-E6 cells (ATCC® CRL-1586™, clone E6) were maintained in Dulbecco’s Modified Eagle Medium (DMEM) complemented with 10% heat-inactivated Fetal Bovine Serum (HI-FBS; PEAK serum), penicillin-streptomycin (Gibco; 15140-122), HEPES buffer (Gibco; 15630-080) and MEM non-essential amino-acids (Gibco; 25025CL) at 37°C with 5% CO2. Vero-CCL81 (ATCC® CRL-81™) cells and MDCK cells (ATCC® CCL-34) were cultured in DMEM supplemented with 10% HI-FBS and penicillin/streptomycin at −37°C with 5% CO2. All experiments involving SARS-CoV2 (USA-WA1/202, BEI resource – NR52281) were conducted within a biosafety-level 3 (BSL3) containment facility at Icahn school of medicine at Mount Sinai by trained workers upon authorization of protocols by a biosafety committee. Amplification of SARS-CoV-2 viral stocks was done in Vero-E6 cell confluent monolayers by using an infection medium composed of DMEM supplemented with 2% HI-FBS, Non-essential amino acids (NEAA), Hepes and penicillin-streptomycin at 37°C with 5% CO2 for 72 hours. Influenza A virus used here was generated using plasmid based reverse genetics system as previously described^34^. The backbone used in the study was A/Puerto Rico/8/34/Mount Sinai(H1N1) under the GenBank accession number AF389122. IAV-PR8 virus was grown and titrated in MDCK as previously described^34^. As a non-enveloped virus, the cell culture adapted murine Encephalomyocarditis virus (EMCV; ATCC® VR-12B) was propagated and titrated in Vero-CCL81 cells with DMEM and 2% HI-FBS and penicillin-streptomycin at 37°C with 5% CO2 for 48 hours^35^.

### 405nm inactivation of viruses

The SARS-CoV-2 virus was exclusively handled at the Icahn school of Medicine BSL-3 and studies involving IAV and EMCV were handled in BSL-2 conditions. Indicated PFU amounts were mixed with sterile 1X PBS and were irradiated in 96 well format cell culture plates in triplicates. In these studies, A starting dose of 5×10^5^ PFU for SARS-CoV-2 and starting doses of 1×10^5^ PFU for IAV and EMCV were used. The final volumes for inactivation were 250 μl per replicate. The untreated samples were prepared the same way and were left inside the biosafety cabinet isolated from the inactivation device at room temperature. The plates were sealed with qPCR plate transparent seal and an approximate 10% reduction of the intensity was observed due to the sealing film. The distance from the lamp and the samples was measured to be 10”. All samples were extracted at indicated times and were frozen at −80°C and were thawed together for titration via plaque assays.

### Plaque assays

For SARS-CoV-2 studies, confluent monolayers of Vero-E6 cells in 12-well plate format were infected (with an inoculum volume 150μl) with 10-fold serially diluted samples in 1X phosphate-buffered saline (PBS) supplemented with bovine serum albumin (BSA) and penicillin-streptomycin for an hour while gently shaking the plates every 15 minutes. Afterwards, the inoculum was removed, and the cells were incubated with an overlay composed of MEM with 2% FBS and 0.05% Oxoid agar for 72 hours at 37°C with 5% CO2. The plates were subsequently fixed using 10% formaldehyde overnight and the formaldehyde was removed along with the overlay. Fixed monolayers were blocked with 5% milk in Tris-buffered saline with 0.1% tween-20 (TBS-T) for an hour. Afterwards, plates were immunostained using a monoclonal antibody against SARS-CoV2 nucleoprotein (Creative-Biolabs; NP1C7C7) at a dilution of 1:1000 followed by 1:5000 anti-mouse IgG monoclonal antibody and was developed using KPL TrueBlue peroxidase substrate for 10 minutes (Seracare; 5510-0030). After washing the plates with distilled water, the number of a plaques were counted. Plaque assays for IAV and EMCV were done in a similar fashion. For IAV, confluent monolayers of MDCK cells supplemented with MEM-based overlay with TPCK-treated trypsin was used and was incubated for 48 hours at 37°C with 5% CO_2_. For EMCV, Vero-CCL81 cells were used to do plaque assays in 6 well plate format with an inoculum volume of 200μl and was incubated for 48 hours at 37°C with 5% CO_2_. Plaques for IAV and EMCV were visualized using crystal violet. Data shown here is derived from three independent experimental setups.

## Results

### Dose and time dependent inactivation of SARS-CoV-2 in the absence of photosensitizers

The lowest irradiation dose of 0.035 mWcm^−2^ was applied for SARS-CoV-2 and when compared to the T_4H_ untreated control, a reduction of 0.3288 log_10_ was seen as early as 4 hours and after 24 hours of irradiation, an inactivation of 1.0325 log_10_ (approximately 10 times reduction in infectivity) was observed for SARS-CoV-2 via plaque assays (Figure 5A). A slightly higher dose of 0.076mWcm^−2^ yielded 0.4123 log_10_, 0.6118 log_10_ and 1.5393 log_10_ reduction by 4, 8 and 12 hours post irradiation when compared to the respective untreated controls (Figure 5B). Subsequent increase of the irradiation dose to 0.150 mWcm^−2^ resulted in a reduction of 0.4771 log_10_ after 4 hours which then had a 1.1206 log_10_ after 12 hours. Irradiation for 24 hours at 0.150 mWcm^−2^ suggested a total reduction of 2.0056 log_10_ (256 times) for SARS-CoV-2 and (Figure 5C). As a final experiment, a high irradiation dose of 0.6 mWcm^−2^ was used to assess the inactivation potential within a much shorter time frame. Irradiation for one hour resulted in a reduction of 0.4150 log_10_ which reached 1.2943 log_10_ reduction by four hours and 2.309 log_10_ (385 times) after 8 hours in comparison to untreated controls samples at the corresponding times. (Figure 5 D and E). All experimental conditions demonstrated the stability of untreated SARS-CoV-2 which was left at room temperature in PBS, as shown by the marginal reduction of viral titer over time.

**Figure 5.**
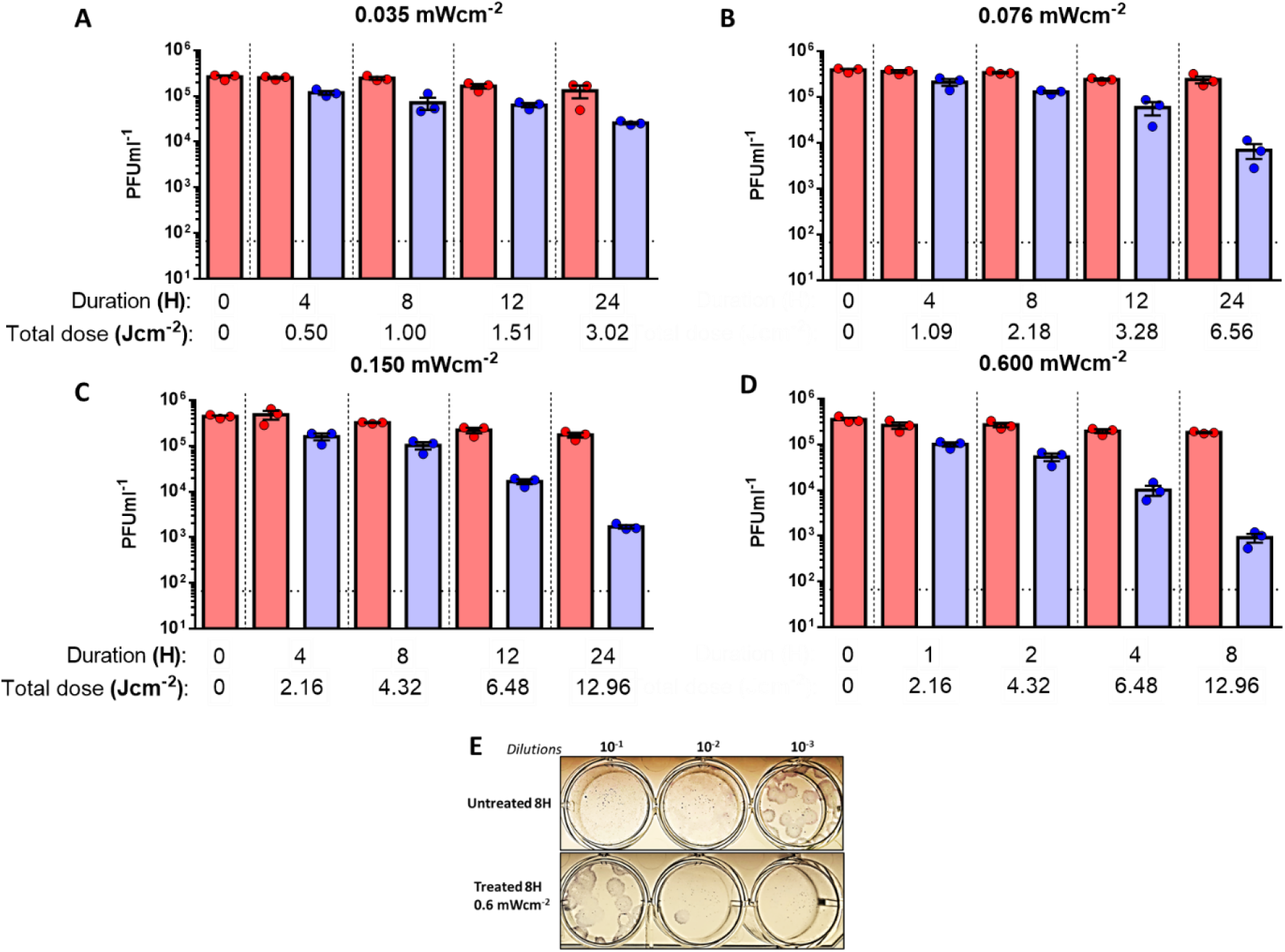
Dose and time dependent inactivation of SARS-CoV-2 virus in PBS by 405 nm irradiation. **A.** A dose of 0.035 mWcm^−2^ or **B.** a dose of 0.076 mWcm^−2^ or **C.**a dose of 0.150 mWcm^−2^ or **D.** a dose of 0.6 mWcm^−2^ was applied to irradiate samples at 405 nm over a course of 24 while sampling at 4, 8, 12 and 24 hours (for A, B and C) or over a course of 8 hours while sampling at 1, 2, 4 and 8 hours (D) was done in independent triplicates. Blue bars indicate treated samples and red bars correspond to the untreated equivalent that was left at the biosafety cabinet under the same conditions while not subjecting to disinfecting irradiation. Data shown as PFUml^−1^ in triplicate assessed by plaque assay. **E.** Plaque phenotype comparison from one independent experiment at an irradiation dose of 0.6 mWcm^−2^. Fixed and blocked plaques were immunostained using anti-SARS-CoV-2/NP antibody before developing using TrueBlue reagent. Data show in here are from three independent replicates (Mean+SEM).

### Influenza A virus is susceptible to 405 nm inactivation in the absence of photosensitizers

Given the observations derived from SARS-CoV-2, a separate inactivation study using a different lipid-enveloped RNA virus was conducted by using influenza A Puerto Rico (A/H1N1/PR8-Mount Sinai) virus strain. Irradiation with a high dose of 0.6 mWcm^−2^ suggested a time dependent reduction of infectious titers as calculated by the 0.1619 log_10_, 0.5609 log_10_, and 1.6115 log_10_ (66 times) reductions at 1, 2, 4 and 8 hours respectively (Figure 6A). And the reduction of plaques was apparent (figure 6B).

**Figure 6.**
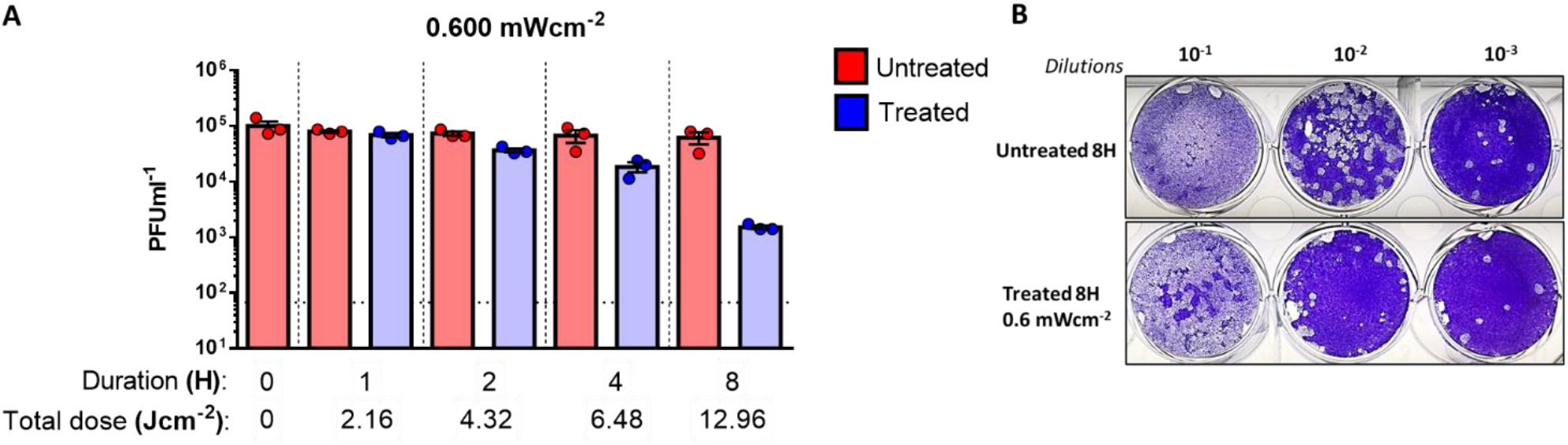
Inactivation of Influenza A virus in PBS by 405 nm irradiation. **A.** A dose of 0.6 mWcm^−2^ was applied to irradiate samples at 405 nm over a course 8 hours while sampling at 1, 2, 4 and 8 hours (done in independent triplicates). Blue bars indicate treated samples and red bars correspond to the untreated equivalent that was left at the biosafety cabinet under the same conditions while not subjected to disinfecting irradiation. Data shown as PFUml^−1^ in triplicate assessed by plaque assay. **B.** Plaque phenotype comparison from one independent experiment at an irradiation dose of 0.6 mWcm^−2^. Fixed and blocked plaques were stained using crystal violet. Data show in here are from three independent replicates (Mean+SEM).

The stability of IAV virus at room temperature for a period of 8 hours was found to be the negligible in untreated IAV spiked PBS samples (Figure 6A).

### Encephalomyocarditis virus (EMCV) as a model non-enveloped virus indicates reduced susceptibility to 405 nm inactivation in the absence of photosensitizers

In order to better understand the effect of the lipid-envelope in viral inactivation by 405 nm irradiation, we used a non-lipid enveloped RNA virus derived from the *Picornaviridae* family. EMCV virus was irradiated at a high dose of 0.6 mWcm^−2^ similar to SARS-CoV-2 and IAV.

In this case however, a total reduction of 0.0969 (approximately 2 times) in comparison to the untreated after 8 hours of irradiation was observed (Fig 7A and 7B) indicating a lower rate of inactivation (despite identical dosing) in contrast to the lipid-enveloped RNA viruses tested in this study. The plaque reduction at 8 hours did not indicate the same dramatic reduction as observed with the latter studies.

**Figure 7.**
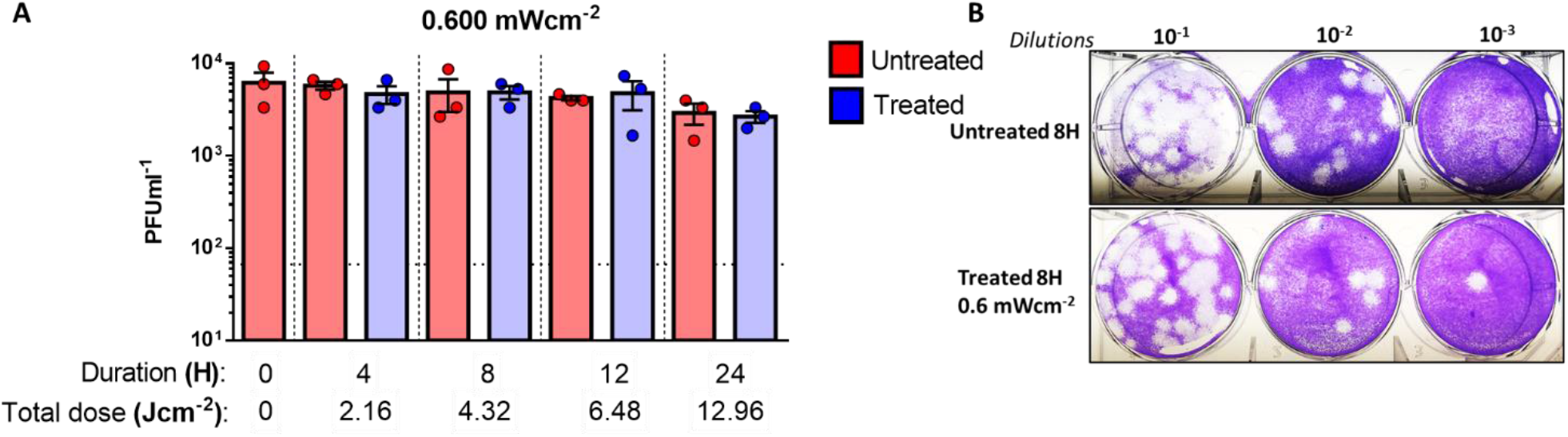
Encephalomyocarditis virus (EMCV) in PBS shows reduced susceptibility to 405 nm irradiation. **A.** A dose of 0.6 mWcm^−2^ was applied to irradiate samples at 405 nm over a course 8 hours while sampling at 1, 2, 4 and 8 hours (done in independent triplicates). Blue bars indicate treated samples and red bars correspond to the untreated equivalent that was left at the biosafety cabinet under the same conditions while not subjected to disinfecting irradiation. Data shown as PFUml^−1^ in triplicate assessed by plaque assay. **B.** Plaque phenotype comparison from one independent experiment at an irradiation dose of 0.6 mWcm^−2^. Fixed and blocked plaques were stained using crystal violet. Data show in here are from three independent replicates (Mean+SEM).

As shown in Figure 4, standard LED lighting (without the specific addition of 405nm light) has a small but measurable amount of disinfecting light (0.006 mWcm^−2^) in the 400nm to 420nm range. To quantify the disinfecting effect of this light, irradiations were performed for 4h, 8h, 12h, and 24 h. A reduction was observed. {Raveen to add detail}. These results at 24h were largely consistent with other irradiation levels (4h, 8h, and 12h) were generally larger than expected even when As shown in Figure 4, standard LED lighting (without the specific addition of 405nm accounting for experimental uncertainty.

## Discussion

The ongoing SARS-CoV-2 pandemic has affected the day-to-day functions in the entire world, raising concerns not only with regards to therapeutics but also in the context of virus survivorship and decontamination^36^. Taking into consideration the rapid spread of SARS-CoV-2 from person to person by droplets, aerosols, and fomites, whole-room disinfection systems can be viewed as a supplement to best practices for interrupting transmission of the virus.

Given the ongoing COVID-19 pandemic, we wanted to explore the impact of 405 nm enriched visible light technology on inactivation of respiratory pathogens such as SARS-CoV-2 and influenza A virus.

Without the use of exogenous photosensitizers, we were able to show that irradiation with low intensity (0.035 mWcm^−2^) visible light yielded a reduction of log_10_ 0.3288 inactivation after four hours (0.5 Jcm^−2^) and a total log_10_ 1.0325 inactivation of SARS-CoV-2 after 24 hours (3.02 Jcm^−2^). A slightly higher dose (0.076 mWcm^−2^) resulted in log_10_ 1.5393 reduction after 24 hours (6.56 Jcm^−2^) while an irradiation dose of 0.150 mWcm^−2^ showed a reduction log_10_ 2.0056 after 24 hours (12.96 Jcm^−2^) of irradiation. Finally, increasing the dose to 0.6 mWcm^−2^ yielded log_10_ 2.3010 reduction only after eight hours (12.96 Jcm^−2^), indicating a both time and dose dependent inactivation of infectious viruses.

The irradiations using standard LED lighting raise some interesting questions for further discussion. With nearly 6x the amount of disinfecting light as compared to traditional fluorescent lighting but 1/6th the lowest amount of 405nm used in the study; it is conceivable that a reduction could be observed. The magnitude of this effect at less than 24h was larger than expected based on other irradiations performed in this study. A similar effect was observed by Bache, et. al {REF} in whole-room bacterial reduction studies. They saw a disinfecting effect for irradiance values as low as 0.023 mWcm^−2^. This effect was shown to be uncorrelated to the level of irradiance used suggesting that the mere abundance of 405nm light can initiate the oxidizing chemical reaction. Nevertheless, our results clearly show a dose-dependent effect. One possible explanation for this observation is that the lower irradiance levels expose different responses by specific organisms within a population based on the individual biology of that organism. Clearly, these results suggest that additional factors may be at work and warrant further investigation.

We selected conventional plaque assays as the read out to specifically estimate infectious virus titers upon disinfection. Methods based in the quantification of viral RNA via PCR based techniques might be misleading as they detect viral RNA from both infectious and noninfectious virions.

SARS-CoV-2 is a lipid-enveloped virus composed of a ssRNA genome and our data indicates its susceptibility to visible light mediated inactivation. To further confirm these observations, we used influenza A virus. which is another human respiratory virus with a lipid envelop and an segmented-RNA genome. Upon irradiating for 1 hour at 0.6 mWcm^−2^ (2.16 Jcm^−2^), we observed a total reduction of log_10_ 0.1619 for the influenza A virus compared to the reduction of log_10_ 0.4150 for SARS-CoV-2 under the same conditions. While both viruses have lipid envelopes, there is clearly a difference here that will require further study. One possible explanation is the difference in the virion size creating a physically smaller cross-section for absorption. (IAV ~120 nm and SARS-CoV-2 ~200 nm)^37, 38^. Nevertheless, both viruses were largely inactivated after eight hours achieving more 1.5 log_10_ reduction. Intriguingly, it was observed that both RNA viruses were able to remain stable at room temperature for at least 24 hours, indicating minimal decay which is consistent with previous studies^36, 39^. We next irradiated a non-enveloped RNA virus, EMCV. Previous results for visible light against non-enveloped viruses demonstrated the need for external photosensitizers such as artificial saliva, blood, feces, etc^30, 36^. Without a porphyrin containing medium, we expected little to no inactivation when this virus was irradiated with visible light. For these measurements, we used the highest available irradiance of 0.6 mWcm^−2^. As anticipated, we observed only a log_10_ 0.0969 reduction after eight hours, however, this appears to be with the statistical precision of the measurement based on the results obtained from shorter irradiations (1, 2, and 4 hours). For comparison, a study involving the M13-bacteriophage virus (a non-enveloped virus) showed a 3-Log reduction using an irradiance of 50mWcm^−2^ (almost 100 times greater than the highest irradiance used in this study) for 10 hours at 425 nm further supporting the idea that non-enveloped viruses may require higher doses of visible light^40^.

Our study was conducted using a neutral liquid media composed of PBS without any photosensitizers and we were able to show that visible light can indeed inactivate lipid-enveloped viruses, differing from the theory that states that photosensitizers are a requirement for inactivation. While these results provide insight into the basic science involved, they were performed within the context of the applied science needed to show the potential impact of this technology upon the current COVID-19 pandemic. By using safe, commercially practical irradiance levels, our results are more directly translatable to occupied rooms in the clinical environment.

Other studies which used visible light-based irradiation have shown similar results in the absence of photosensitizers, indicating the possibility of an alternative inactivation mechanism^23, 25, 30^. Studies have proposed two theories for this observation primarily due to non-405nm wavelengths emitted by the source: 1) some amount of 420-430 nm emitted from the source is contributing to the viral inactivation ^41^, and 2) the presence of UV-A (390 nm) within the source. This wavelength is known to create oxidative stress upon viral capsids^42^.

Longer wavelengths, such as 420-430nm, have shown inactivation of the murine leukemia virus (MRV-A) ^41^. While this is an intriguing study, it used a broad-spectrum lamp with optical filters to selectively identify the spectrum primarily responsible with their results. Unfortunately, they did not quantify the amount of light (using radiometric units) within the spectrum of interest used to irradiate the virus. While transmission profile of the filters used were provided, it does not consider the spectral composition of the source itself making any direct quantitative comparison between our studies impossible. It is interesting to note that they did observe viral inactivation in their controls from wavelengths less than 420nm confirming the qualitative findings of our study without confirming the specific use of 405nm. This suggests that the viral inactivation is a likely a broad response (> 20nm) with relative contributions unique to the chemistry of each organism. They also considered much longer exposures (~7 days) and much higher illuminance (> 200 lux) than that used in our study although this is again difficult to compare given the lack of radiometric quantification of their light source. It is important to note that the control samples used in our study were exposed to the same overhead (non-405nm) lights as the irradiated samples and our results are the observed difference between the two demonstrating the contribution from 405nm over and above that potentially from 420-430nm. Future experiments can further quantify the potential effect.

The other theory, potential UV-A irradiation, was historically applied to lamp-based sources with broad spectral (> 100nm) outputs. Again, the use of LED technology addresses this question as the peak irradiance at 390nm of the device used in this study was < 1% of its peak irradiance at 405nm without the need for any additional filtration. Future experiments can further quantify the potential effect.

Another consideration to be addressed is thermal heating of the virus by the LED source. Tsen and Achilefu used a pulsed laser method at 425nm^43^ with ~100 mWcm^−2^ average power density for < 2h while simultaneously measuring the sample temperature with a thermocouple. They detected less than a 2°C demonstrating minimal temperature impact even under a power density nearly 9 orders of magnitude larger than that used in this study. This was confirmed by our thermocouple measurements as stated earlier. Nazari, *et al* used an 805nm source with an average power density of > 0.3 Wcm^−2^ for 10s, nearly 1000 times that used in this study^44^. While the total energy delivered was more comparable to that used in our study, they did not make explicit temperature measurements, their analysis ruled out any potential thermal effects.

One possible explanation for the observed differences between the enveloped and non-enveloped organisms is absorption of the 405nm light by the lipid envelope itself. This could, in turn, lead to the creation of reactive oxygen species (causing an oxidative effect) or simply destruction of the envelope leading to a denaturing of the organism. This question could serve as the basis for a range of future studies.

The results obtained suggest that the performance of visible light against SARS-CoV-2 is similar to other organisms commonly found in the environment such as *S. aureus*. Previous studies have shown that the visible light irradiance levels used in this study (0.035 mWcm^−2^ to 0.6 mWcm^−2^) reduce bacteria levels in occupied rooms and improve outcomes for surgical procedures. It is therefore reasonable to conclude that visible light might be an effective disinfectant against SARS-CoV-2. More importantly, this disinfection can operate continuously as it is safe for humans based upon the exposure guidelines in IEC 62471^45^. This means that once it has been in use for a period of time, the environment will be cleaner and safer the next time it is occupied by humans.

One limitation of this study is that the inactivation assays were performed in static liquid media as opposed to aerosolized droplets. While the use of visible light in air disinfection has been briefly studied where it was shown that its effectiveness increased approximately 4-fold^46^, further studies involving dynamic aerosolization are needed to better understand the true potential of visible light mediated viral inactivation.

In any case, our study shows the increased susceptibility of enveloped respiratory viral pathogens to 405 nm mediated inactivation in the absence of photosensitizers. The irradiances used in this study are very low and might be easily applied to disinfect occupied areas safely and continuously within hospitals, schools, restaurants, offices and other locations. Of particular interest is the potential for standard LED lighting to play a role in reducing the presence of SARS-CoV-2 in the environment.

## Conclusions

We have demonstrated the basic science of inactivation of enveloped viruses such as SARS-CoV-2 and Influenza-A using 405nm visible light within the context of the applied science required for this technology to have an impact upon the current COVID-19 pandemic. Without the need for exogenous photosensitizers and by using safe, commercially practical irradiance levels, our results can be easily translated to the clinical environment.

### Future Efforts

Future work should focus on explaining the difference between the enveloped and non-enveloped results. This may include transmission electron microscopy (TEM), hemagglutination assay (HA), or other methods focusing on the potential role that a mediated reaction (due to the envelope itself) might play. The size of the virion particle may play a role in photoelectric absorption and could be studied for different viral species. We acknowledge that while unlikely, other wavelengths of visible light, beyond 400-420nm may play a role in the inactivation process and future studies should explore this possibility as well. Finally, the inactivation kinetics of low irradiances could add valuable insight into clinical applications of this technology.

## Acknowledgments

We thank Kenall Manufacturing for supplying the M4DLIC6 fixture and test rig used in this study. We thank Randy Albrecht for support with the BSL3 facility and procedures at the ISMMS, and Richard Cadagan for technical assistance. This research was partly funded by CRIP (Center for Research for Influenza Pathogenesis), a NIAID supported Center of Excellence for Influenza Research and Surveillance (CEIRS, contract # HHSN272201400008C); by the generous support of the JPB Foundation, the Open Philanthropy Project (research grant 2020-215611 (5384)) and anonymous donors to AG-S, and by a research contract from Kenall Manufacturing to the AG-S lab.

